# High-Resolution Metabolic Mapping of the Cerebellum Using a Zoomed Magnetic Resonance Spectroscopic Imaging

**DOI:** 10.1101/2020.05.20.093393

**Authors:** Uzay E. Emir, Jaiyta Sood, Mark Chiew, Albert Thomas, Sean P. Lane

## Abstract

**Purpose:** The human cerebellum plays an important role in functional activity cerebrum which is ranging from motor to cognitive activities since due to its relaying role between spinal cord and cerebrum. The cerebellum poses many challenges to magnetic resonance spectroscopic imaging (MRSI) due to the caudal location, the susceptibility to physiological artifacts and partial volume artifact due to its complex anatomical structure. Thus, in present study, we propose a high-resolution MRSI acquisition scheme for the cerebellum.

**Methods:** A zoomed or reduced-field of view (rFOV) metabolite-cycled full-intensity magnetic resonance spectroscopic imaging (MRSI) at 3T with a nominal resolution of 62.5 μL was developed. Single-slice rFOV MRSI data were acquired from the cerebellum of 5 healthy subjects with a nominal resolution of 2.5□×□2.5□mm2 in 9□minutes 36. Spectra were quantified with LCModel. A spatially unbiased atlas template of the cerebellum was used for analyzing metabolite distributions in the cerebellum.

**Results:** The high quality of the achieved spectra enabled to generate a high-resolution metabolic map of total N-acetylaspartate, total creatine, total choline, glutamate+glutamine and myo-inositol with Cramér-Rao lower bounds below 50%. A spatially unbiased atlas template of the cerebellum-based region of interest (ROIs) analysis resulted in spatially dependent metabolite distributions in 9 ROIs. The group-averaging across subjects in the Montreal Neurological Institute-152 template space allowed to generate a very high-resolution metabolite maps in the cerebellum.

**Conclusion:** These findings indicate that very high-resolution metabolite probing of cerebellum is feasible using rFOV or zoomed MRSI at 3T.

## Introduction

Compared to the cerebrum, cerebellum has only 10% of total brain volume, however it has 3 times more neurons than the cerebrum (1) and consumes around 20% of the whole energy utilization of the brain (2). The human cerebellum plays an important role in functional activity of the brain which is ranging from motor to cognitive activities due to its relaying role between spinal cord and cerebrum. With the development of the neuroimaging modalities, the cerebellum has been the focus of interest but its complex structure and caudal location inherits technical challenges for imaging modalities (3). Increasingly, advanced MRI technologies including anatomical and functional modalities have facilitated the evaluation of the cerebellum (4,5). This is of great importance since abnormalities in the cerebellum are associated with many neurological, neurodevelopmental, and psychiatric disorders (6–9).

Magnetic resonance spectroscopy (MRS) techniques are increasingly used for to evaluate the neurochemical probing of the brain tissue (10). Single voxel (SV) MRS has been the method of choice to study the cerebellum with a nominal voxel dimension ranging from 4090 μL (11) to 15620 μL (12) for different neurological conditions. However, spectra in single SV-MRS acquisitions are derived from a mixture of different tissue compartments, which results in a partial volume effect. This partial volume effect exacerbates in the cerebellum due to its complex structures.

Alternatively, MRS imaging (MRSI) methods record multiple spectra from different regions simultaneously. However, the human cerebellum poses many challenges to MRSI research due to the caudal location and the susceptibility to physiological artefacts. Only very few studies have been performed using ^1^H MRSI in the cerebellum with coarse nominal resolutions ranging from 310 μL (13) to 1210 μL (14).

Zoomed-MRI or reduced-field of view (rFOV) acquisition schemes overcome the limitations of spatial resolution and low signal to noise ratio (SNR) for MRI. This has been achieved by using an outer volume suppression (OVS) (15) or a well-defined spatial excitation (16). Similar to the Zoomed MRI, three-pulse localization of MRSI suppresses the signal outside of the FOV and used for rFOV MRSI measurements (17,18). The present study, therefore, sought to develop an accelerated acquisition scheme of rFOV for high-resolution full-intensity short echo-time (TE) semi-LASER MRSI at 3T clinical scanners with a nominal resolution of 62.5 μL. In five healthy subjects, we demonstrate that the proposed rFOV-MRSI with accelerated k-space trajectory generates high-quality metabolic maps at 3 Tesla across an entire cerebellum slice with a thickness of 10 mm. Finally, we explored neurochemical alterations in regions of interests (ROIs) of the cerebellum using the spatially unbiased atlas template of the cerebellum (SUIT Atlas) (4).

## Methods

Five healthy subjects [three females, 25.5□±□7.07 years (mean□± □std)] participated. All subjects provided informed consent before the in-vivo MRI exam which was approved by the Purdue University institutional review board.

### MRI data acquisition

The data were acquired using Siemens Prisma 3T MR system (Siemens, Germany) with a 64-channel (N_channels_) head array receive coil. A T1-weighted MP-RAGE dataset (TR□=□1900 ms, TE□=□3.97□ms, TI□=□904□ms, flip angle□=□8°, 192 transverse slices, 1□mm^3^ isotropic voxels) was acquired for each subject for MRSI acquisition planning. B0 shimming was achieved vendor-provided procedure, GRESHIM (gradient-echo shimming).

### Reduced field of view, Zoomed, metabolite-cycled semi-laser MRSI

Metabolite-cycling MRSI was acquired using the same parameters as described in Emir et al. (19). Briefly, before the semi-LASER localization, two asymmetric inversion RF pulses in alternating TRs were used for downfield/upfield (N_directions_ = 2 [upfield/downfield]) measurements.

The semi-LASER localization box encompassing the entire cerebellum (Figure 1A) was positioned to include Right Crus I, Left Crus I, and Dentate Nuclei (Figure 1B). The high in-plane nominal resolution (2.5 mm × 2.5 mm) with a thickness 10 mm was achieved using rFOV = 120 mm × 120 mm × 10 mm, semi-LASER localization = 100-90 mm × 50-40 mm × 10 mm, TR = 1500 ms, and TE = 32 ms.

**Figure 1.**
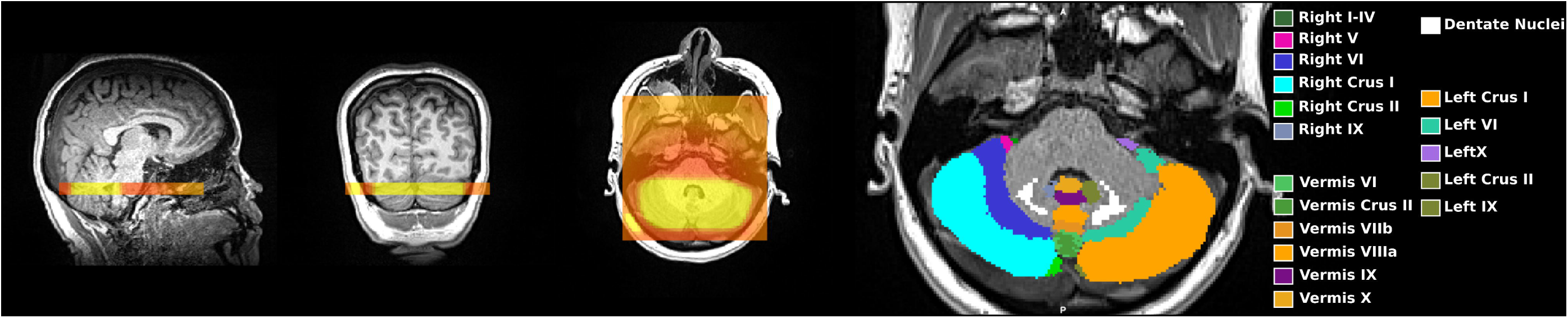
(A) Representative placement of the MRSI slice on MP-RAGE images. (B) Typical region of interest (ROI) analysis is illustrated on the same subjects’ MP-RAGE axial image.

To cover the 48 × 48 grid in a rFOV, we have modified our 2D-density weighted concentric ring trajectory (DW-CRT) design (20). Each ring in DW-CRT consisting of 64 points (number of points per ring, N_p_ring_ = 64) was collected with an ADC bandwidth of 40 kHz. Two temporal interleaves (number of temporal interleaves, N_ti_ = 2) with inverted readout gradients were collected with an SBW of 625 Hz (20). This resulted in 128 spectral points (N_sp_ = 128). The rest of the DW-CRT parameters were identical to the previous 3T and 7T implementations (20,21): four spatial interleaves (N_si_ = 4 and N_avg_ = 1) of 24 DW concentric rings (N_ring_ = 24) were acquired with α = 1. The total duration for the DW-CRT was 9 minutes 36 seconds ((N_si_ = 4) x (N_directions_□=□2) × (N_ring_□=□24) x (N_ti_ = 2) x (TR = 1500 ms) = 576 seconds).

### Post processing

MATLAB (MathWorks, Natick, MA, USA) was used for reconstruction algorithms. To achieve the full SBW (1250 Hz), the NUFFT (22) was used to calculate the Fourier transform of the acquired k-space data (20).

Single-shot metabolite-cycled spectra were processed as described in Emir, et al. (19). The non-water-suppressed spectra enabled to perform frequency and phase corrections for each voxel with a dimension of 62.5 μL. A Gaussian filter of 250-ms timing parameter were applied to FIDs. The metabolite spectra were obtained via subtracting the upfield and downfield FIDs while summing them generated the water-only spectra. The remaining water signal in the metabolite spectrum was removed using the Hankel-Lanczos singular value decomposition (23). The lipid-basis penalty algorithm was used to minimize the lipid contamination (24). The water-only and metabolite-only whole cerebellum slice images were used to generate a brain and lipid mask for the lipid-basis penalty algorithm.

### Metabolite Quantification

Metabolite quantification was performed using the LCModel (25). The model spectra including eight LCModel-simulated macromolecule resonances was used as a basis-set for LCModel analysis, as described in Emir, et al. (21). The resulted Cramér-Rao lower bounds of estimated metabolites less than 50 % were reported. Summed metabolite estimations were reported for total N-acetylaspartate (NAA+NAAG, tNAA), total creatine (Cr + PCr, tCr), total choline (GPC + PC, tCho) and glutamate and glutamine (Glu + Gln, Glx), since their pairwise-correlation estimates were high (correlation coefficient < −0.5). The normalized metabolite levels were reported for each subject using the water-scaled signal intensity of each metabolite (raw): Metabolite-normalized=Metabolite-raw/(tCho_raw_+tCr_raw_+tNAA_raw_+*myo*-Ins_raw_+Glx_raw_).

### Regional Distributions of Neurochemical in the Cerebellum

The Cerebellar ROIs were obtained using the Cerebellar MNI FNIRT Maxprob thr25-2mm of SUIT Atlas (http://www.diedrichsenlab.org/imaging/propatlas.htm). Each subject’s MRSI slice was determined in the MP-RAGE image space using SPM (26). Then, MRSI metabolite maps for each subject warped in the Montreal Neurological Institute (MNI)-152 template in the following manner, each subject’s T1 image was nonlinearly registered to the Montreal Neurological Institute-152 template by using the FMRIB Nonlinear Registration Tool (FNIRT); afterwards, the resulting warp field was applied to the Cerebellar MNI FNIRT Maxprob thr25-2mm of SUIT atlas to transform them to each subjects MPRAGE image. Mean concentration in cerebellar ROIs were calculated using fslstats (27).

To generate group-averaged metabolite spatial maps, metabolite maps were transformed to the MNI-152 template by applying the resulting warp field; afterwards metabolite maps were averaged across five subjects.

### Statistical Analysis

To select reliable ROIs, ROIs of the cerebellar atlas encompassed by at least the half of the subjects’ MRSI box were reported. As a secondary filter to select reliable ROIs, the mean number of voxels in a ROI across subjects had to be one standard deviation higher than the standard deviation of the number of voxels in an ROI.

The mean regional Metabolite-normalized levels from the aforementioned reliable ROIs of cerebellar atlas are used for statistical analyses using SAS software v.9.4 (SAS Institute, Cary, NC, 2014). MRS data from different ROIs were compared using a one-way repeated measures analysis of variance separately for each reported normalized metabolite (tNAA, tCr, tCho, Glx and *myo*-Ins) across each target brain region. Given the occurrence of missing data due to exclusion for unreliability, we utilized restricted maximum likelihood to fit the models. Each analysis examined the differences in MRS data across 9 ROIs, resulting in 36 possible pairwise comparisons. We employed a Benjamini-Hochberg correction within each family of comparisons in order to control the false discovery rate (28).

## Results

Figure 1A shows coronal, sagittal, and axial images of an anatomical scan including the placement of the MRSI slice and semi-LASER localization. The image derived from the non-water suppressed MRSI data shows the signals from the cerebellum. ROIs of the SUIT atlas encompassed by the MRSI localization are illustrated in Figure 1B. The list of ROIs and the mean number of voxels for each metabolite that met our criteria for reliable quantification in the SUIT atlas are listed in Table 1.

**Table 1.**
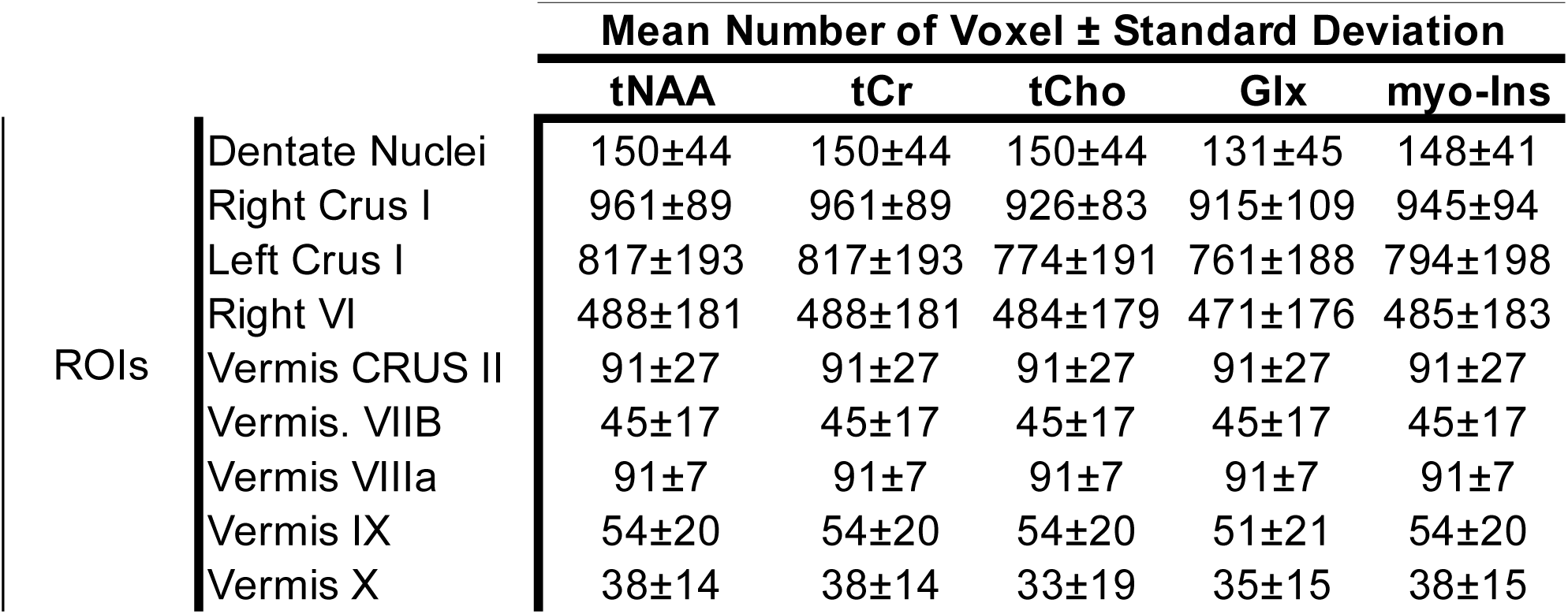
The list of ROIs and the mean number of voxels across subjects for each metabolite. ROIs are in rows and metabolites are in columns.

Figure 2 shows the zoomed MRSI spectra with LCModel fit. Even at a nominal resolution of 62.5 μL at 3T, spectra from five different volumes of interest are of high quality.

**Figure 2.**
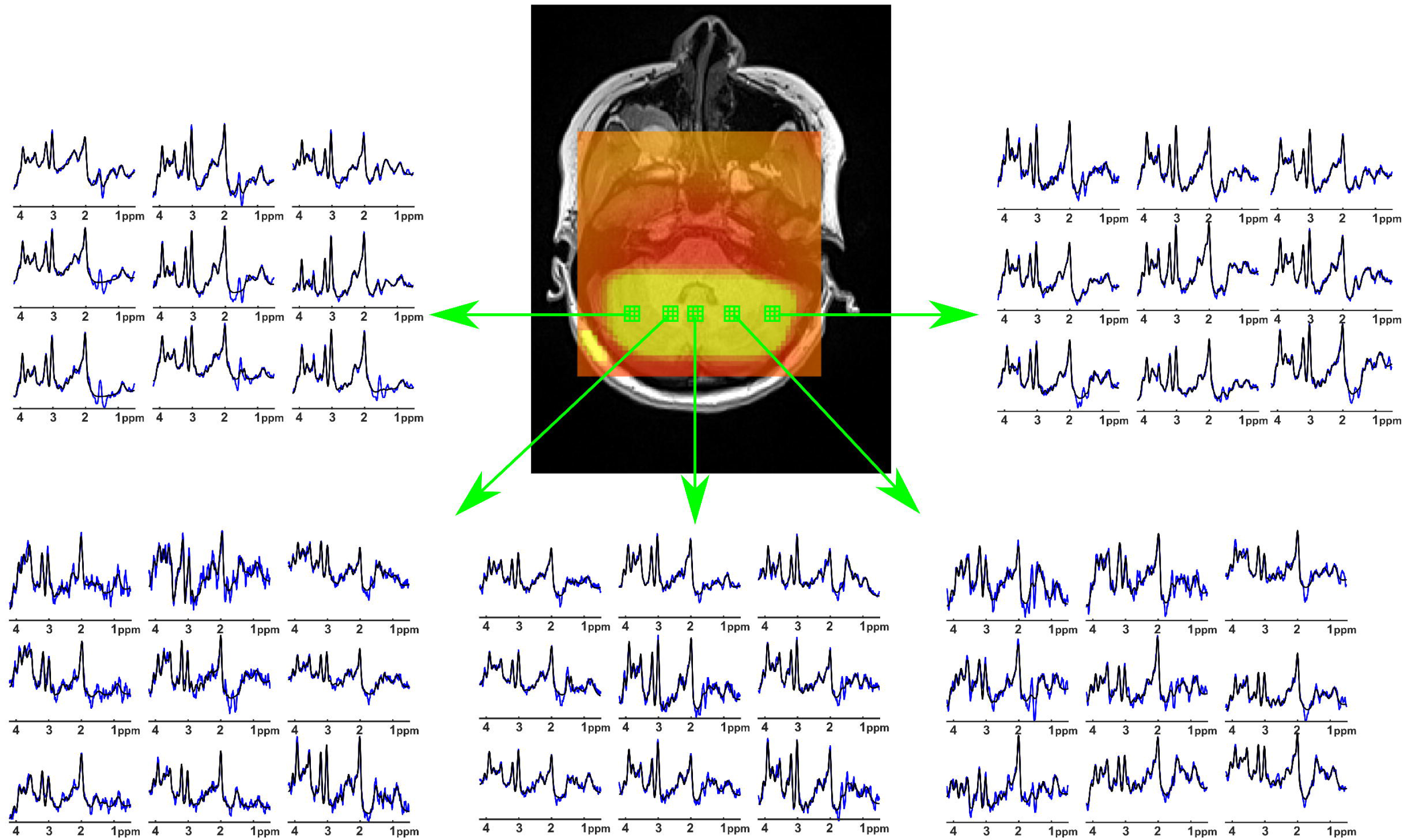
Extracted spectra (blue) from nine voxels (2.5 mm x 2.5 mm x 10 mm each) of 5 different volumes of interests with LCModel fit (black).

In most of the voxels, the CRLB values were resulted in less than 50% for the metabolites of tNAA, tCr, tCho, myo-Ins, and Glx. The resulting normalized metabolite maps and corresponding CRLB maps of a healthy subject are illustrated in Figure 3A and Figure 3B, respectively.

**Figure 3.**
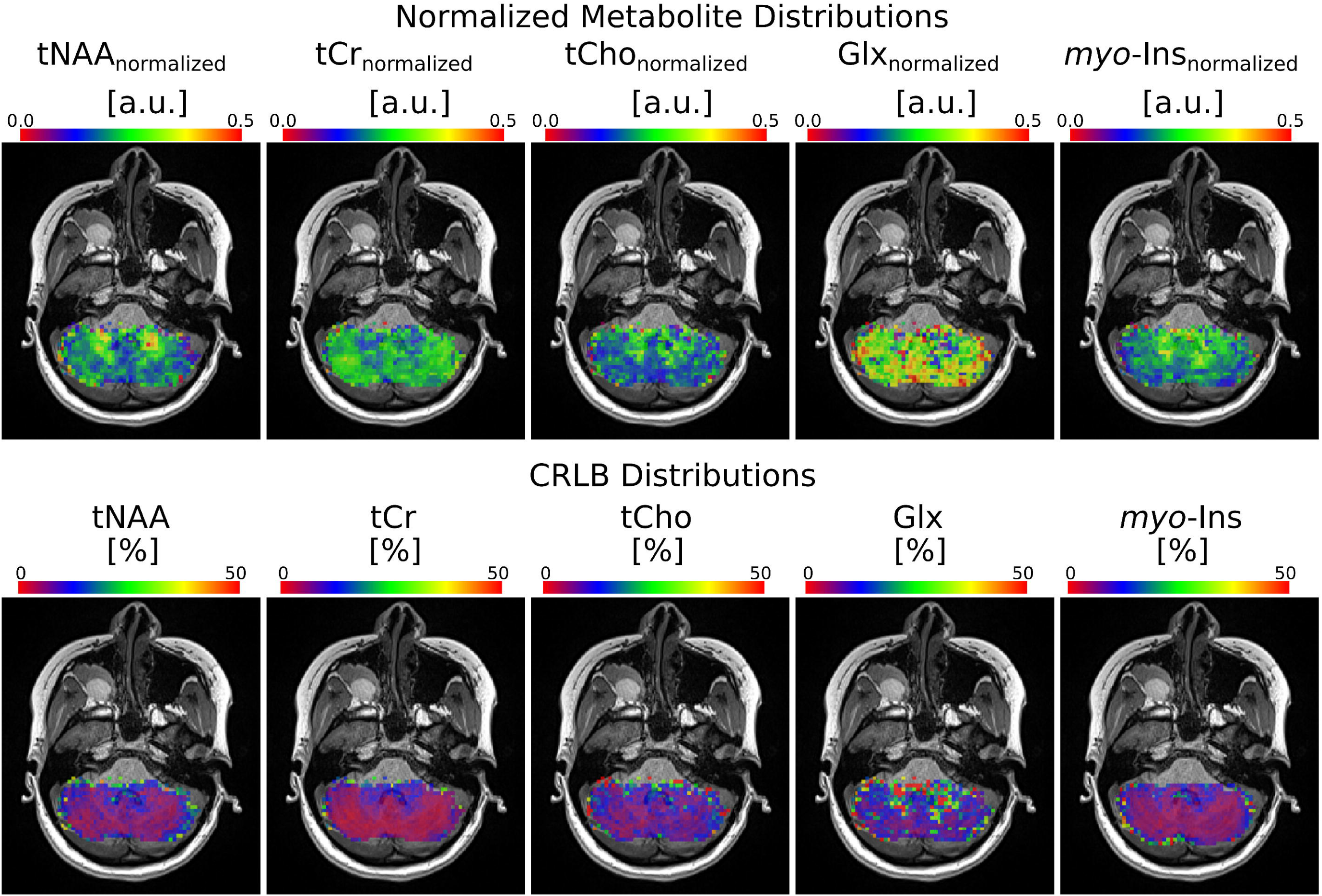
Metabolite and CRLB distribution maps obtained with rFOV MRSI from a subject. Normalized metabolite level maps from a nominal voxel dimension of 62.5 μL) for tNAA, tCr, tCho, Glx and myo-ins overlaid on an anatomical image. a.u.: arbitrary units

Figure 4 shows the group-average metabolite maps in the MNI-152 template space. The mean normalized metabolite levels that met our criteria for reliable quantification in cerebellar ROIs (nine cerebellar ROIs) of SUIT atlas are shown in Figure 5. The comparison of normalized metabolite levels revealed neurochemical profiles characteristic of different cerebellar regions (Table 1S, supporting material). For instance, the tCho, tNAA, and myo-Ins levels in the Dentate Nuclei are significantly higher than the Right and Left Crus I and Vermis ROIs, whereas tCr is significantly lower compared to the Right and Left Crus I and Vermis ROIs. Similarly, Dentate Nuclei has higher tCho, tNAA, and myo-Ins levels compared to the Vermis ROIs.

**Figure 4.**
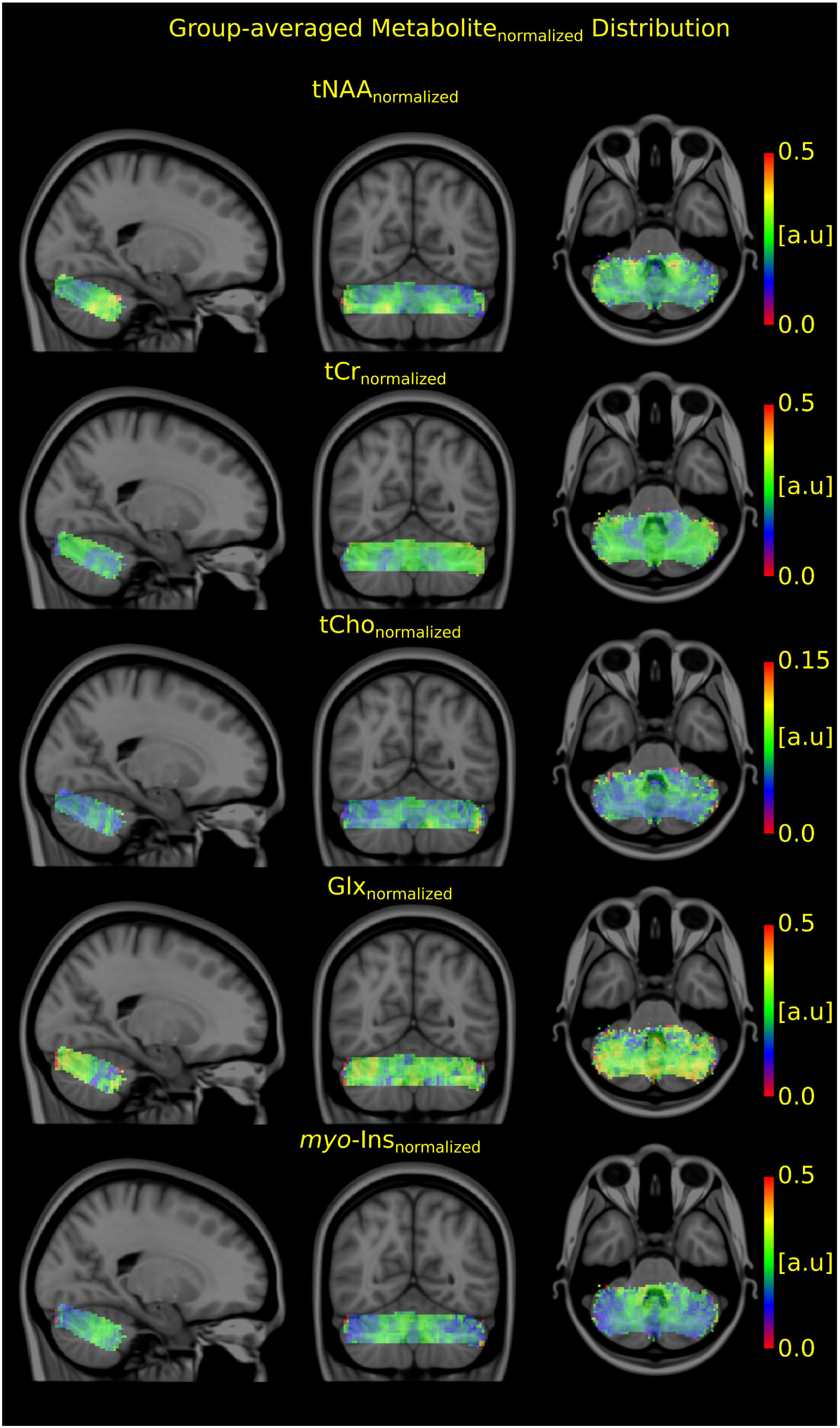
Group-average normalized metabolite maps overlaid on Montreal Neurological Institute-152 template. a.u.: arbitrary units.

**Figure 5.**
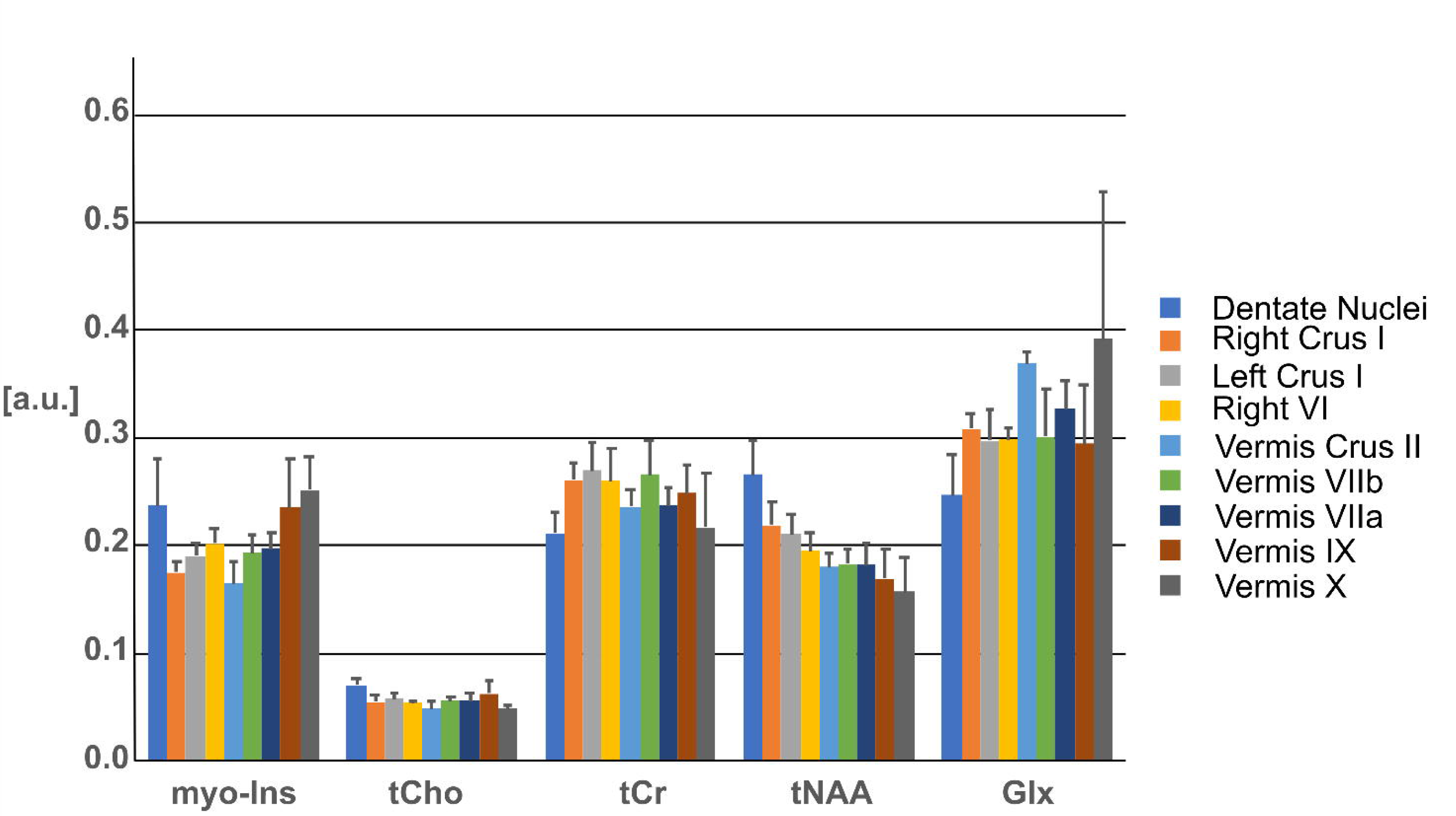
Neurochemical profiles from different brain regions determined by LCModel fitting of rFOV MRSI with a nominal resolution of 2.5 mm × 2.5 mm × 10 mm acquired at 3T. Only metabolites and ROIs that were quantified reliably are shown. Metabolites were significantly different between regions p < 0.05 (ANOVA). Error bars: inter-subject SD (N=5 for all brain regions). a.u.: arbitrary units.

## Discussion

This study has demonstrated the advantage of rFOV acquisition, zoomed, with a DW-CRT short TE semi-laser localization to improve the resolution and accuracy of whole cerebellum slice metabolic maps at 3T. The improved resolution and spectral quality enabled reliable quantification of tNAA, tCr, tCho, Glx and *myo-*Ins from a nominal voxel of 62.5 μL. The metabolite distribution maps for different ROIs in the cerebellum were consistent with the literature (see below). To our knowledge, this is the first study to have validated a higher nominal resolution of 62.5 μL at 3T.

Implementing rFOV acquisition at 3T for high-resolution MRSI faces several challenges caused by the hardware and signal-to-noise ratio limitations. Mainly, high-resolution MRSI puts demands on both the maximum gradient amplitudes, rise time, and slew rate. To decrease the field of view while maintaining the acquisition matrix size, one might choose to increase the gradient amplitude and slew rate or to reduce the spectral bandwidth resulting in lower spectral sampling. In this study to reach the nominal in-plane resolution of 2.5 mm × 2.5 mm, we decided to utilize reduced SBW (625 Hz vs 1250 Hz) compared to the previous implementation (21). The SBW was doubled (1250 Hz) using an inverted readout gradient trajectory without causing any additional noise (20). The use of DW-CRT and its reconstruction pipeline further contributed the SNR improvement as discussed previously in Chiew et al. (20). Besides, a lower acquisition bandwidth of 40 kHz compared to the previous implementation of 80 kHz (19,21,29) minimized a larger amount of noise sampled due to the narrower frequency range. Moreover, the 64-ch Head/Neck coil allowed the incremental SNR gains in the peripheral ROIs closest to the coil elements, like the cerebellum (30). Finally, additional SNR improvement was achieved through the use of metabolite-cycling allowing to conduct voxel-vise preprocessing steps (19).

The present study did not directly compare this acquisition to others at 3Tesla in which the nominal resolution ranged from 300 μL (31) to 1000 μL (32). However, the resulted nominal resolution and metabolite maps achieved in this study are comparable (62.5 μL vs 40 μL) to the ones in which constrained reconstruction methods with FID acquisition were used (33,34). The achieved nominal resolution and provided metabolite maps within 9 minutes 36 seconds are also comparable with the ultra-high-field (UHF ≥ 7T) ones with nominal voxel sizes of 23 μL (35). Thus, we believe that our strategy offers substantial resolution improvement compared to the other implementations at 3T and UHF. With the use of constrained reconstruction methods, the proposed strategy is expected to provide higher resolution (33,34).

The rFOV MRSI acquisition with a resolution of 62.5 μL allowed reliable quantification of major brain metabolites in the cerebellum (Figure 2 and 3). Concentration distributions in the cerebellum of the reported metabolites revealed significant variations between the cerebellar vermis, white, and gray matter of both cerebellar hemispheres and these were inline with previous MRSI and SV-MRS studies. For example, the high cerebellar total creatine level in the gray matter agreed with previous reports (14). In addition, the higher Glx level obtained in cerebellar gray matter was consistent with the previous SV-MRS studies (36). The highest total choline levels were observed in the cerebellar white matter, also in agreement with SV-MRS (36).

The voxel-based comparisons of SUIT Atlas ROI (4) analyses of whole cerebellum slice metabolite maps demonstrates that metabolite distributions are spatially dependent. The dentate nuclei, the link in the cortico-cerebellar close loop circuits (37), had higher tNAA, tCho, and myo-ins levels. Since NAA was extensively distributed in the cerebellum and NAAG was very high in the deep cerebellar nuclei, higher tNAA levels observation in the dentate nuclei is in agreement with previous studies (38). Since the dentate nuclei are embedded in the white matter having a higher glia-to-neuron ratio than grey matter (39), myo-inositol and tCho as markers of glial cells resulted in higher levels in the dentate nuclei. Inline with the cyto and receptor architectonic mapping of glutamate receptors in the human cerebellum (40), Glx level is higher in the cerebellar cortex (Right Crus I, Left Crus I and Vermis) than in the dentate nuclei. Finally, higher creatine levels in the cerebellar cortex might suggest energy demand of Purkinje cells (14).

Qualitatively, the combination of the 2D MRSI acquisitions across subjects improved the metabolite distributions resolution from 62.5 μL to 8 μL since the different MRSI slice placements across subject provide sufficiently different cerebellum coverage. With the accurate coregistration between MRSI and MNI template, the high-quality group-average metabolite distributions of tNAA, tCr, tCho, Glx and myo-Ins were achieved.

The number of subjects is a limitation and further experiments are required for the clinical validity of the proposed method. In addition, single slice acquisition is not sufficient enough to map the entire cerebellum. However, this study has shown that multiple slices with different orientations may provide more reliable metabolite distribution (41). The use of long TR of 1.5 s and the cost of increased total acquisition duration due to the choice of adiabatic pulses can be mitigated by gradient-modulated offset-independent adiabaticity (GOIA) pulses (42). Together with temporal interleaves (N_ti_ = 2) total acquisition was 9 minutes 36 seconds. Recently, we have reduced acquisition duration to 4 minutes 48 seconds by reducing the spectral bandwidth to 893 Hz (N_ti_ = 1) with the use of maximum slew rate of the gradient system (data was not shown). Finally, the exact quantification of peaks might be affected by the use of simulated macromolecules and lipid removal procedure.

In conclusion, the rFOV 2D MRSI resulted in very high-resolution metabolite maps of the cerebellum. These pilot findings indicate optimal methods which can be developed for the future to generate probabilistic metabolic atlas of the cerebellum.

## Abbreviations

2D: two dimensional;
ANOVA: analysis of variance;
CRLB: Cramér-Rao lower bound;
CRT: concentric ring trajectory;
DW: density weighted;
FID: free induction decay;
FOV: field of view;
GABA: γ-aminobutyric acid;
Gln: glutamine;
Glu: glutamate;
GPC: glycerophosphocholine;
GRESHIM: gradient-echo shimming;
GSH: glutathione;
HLSVD: Hankel-Lanczos singular value decomposition;
Lac: lactate;
MNI: Montreal Neurological Institute;
MRSI: magnetic resonance spectroscopic imaging;
myo-Ins: myo-inositol;
NAA: N-acetylaspartate;
NAAG: N-acetylaspartylglutamate;
PCho: phosphocholine;
PCr: phosphocreatine;
rFOV: reduced field of view;
SAR: specific absorption rate;
SBW: spectral bandwidth;
SD: standard deviation;
semi-LASER: semi-localization by adiabatic selective refocusing;
SNR: signal-to-noise ratio;
SUIT: spatially unbiased atlas template;
tCho: total choline;
tCr: total creatine;
VOI: volume of interest

---

**Table 1S.**
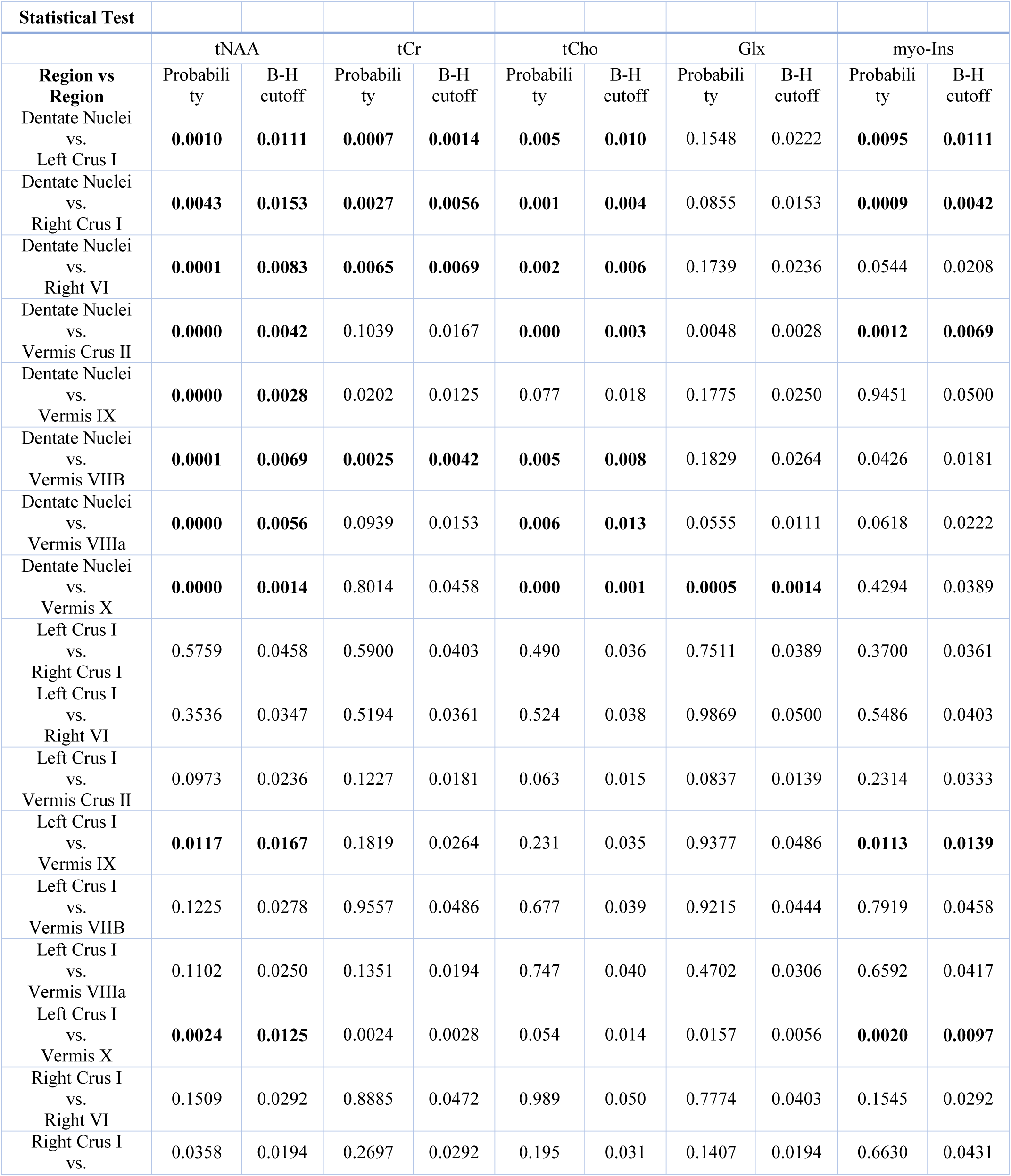
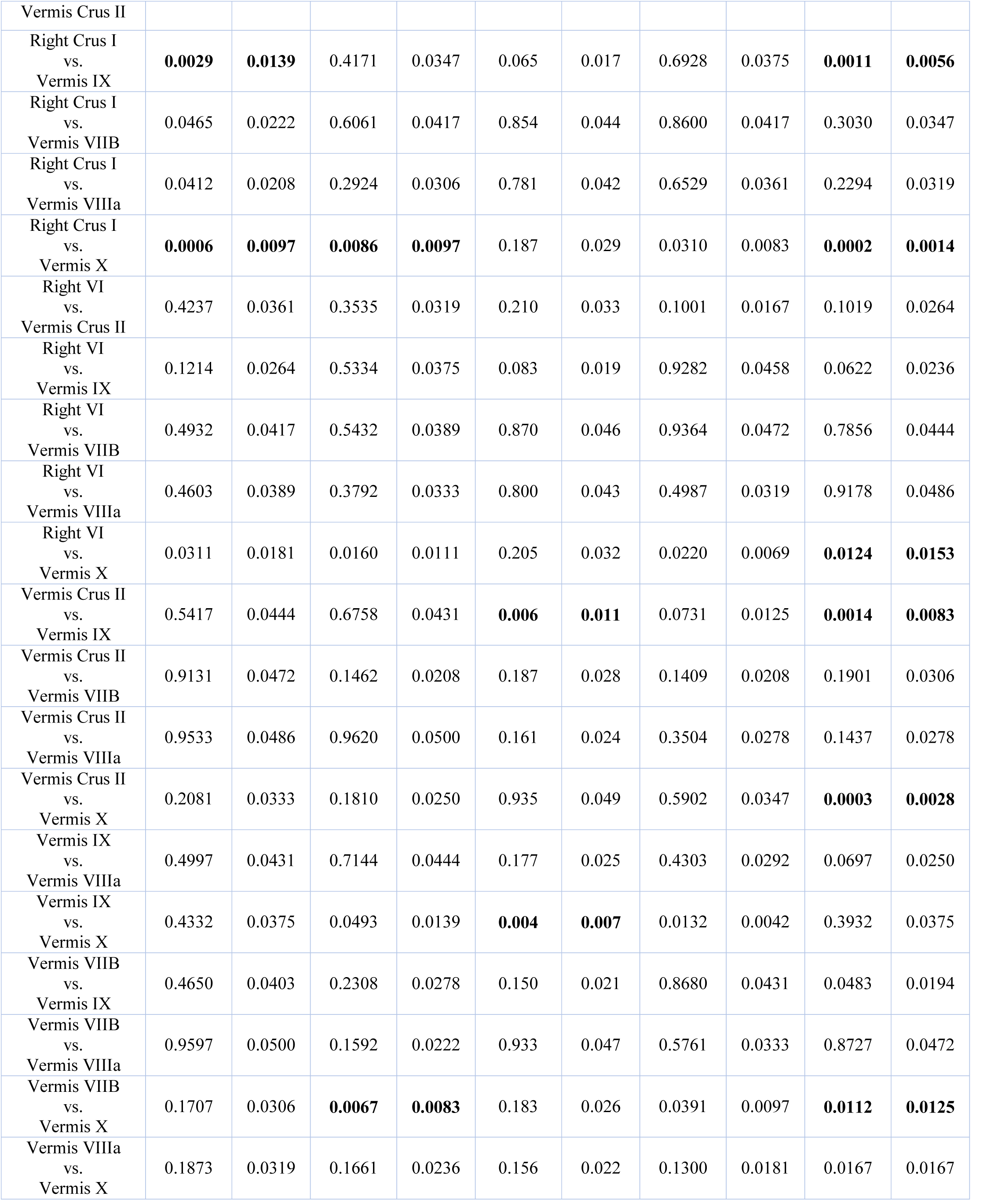
Pairwise contrasts with explicit p-values and Benjamini-Hochberg corrected versions of the differences in MRS data across 9 ROIs, resulting in 36 possible pairwise comparisons. We employed a Benjamini-Hochberg correction (B-H cutoff) within each family of comparisons in order to control the false discovery rate of 5 %. Statistically significant P-values are highlighted in bold font.

## References

1. Van Essen DC, Donahue CJ, Glasser MF. Development and Evolution of Cerebral and Cerebellar Cortex. Brain Behav Evol 2018;91:158–169 doi: 10.1159/000489943.

2. Howarth C, Gleeson P, Attwell D. Updated energy budgets for neural computation in the neocortex and cerebellum. J Cereb Blood Flow Metab 2012;32:1222–1232 doi: 10.1038/jcbfm.2012.35.

3. Diedrichsen J, Verstynen T, Schlerf J, Wiestler T. Advances in functional imaging of the human cerebellum. Curr. Opin. Neurol. 2010;23:382–387 doi: 10.1097/WCO.0b013e32833be837.

4. Diedrichsen J, Balsters JH, Flavell J, Cussans E, Ramnani N. A probabilistic MR atlas of the human cerebellum. Neuroimage 2009;46:39–46 doi: 10.1016/j.neuroimage.2009.01.045.

5. King M, Hernandez-Castillo CR, Poldrack RA, Ivry RB, Diedrichsen J. Functional boundaries in the human cerebellum revealed by a multi-domain task battery. Nature Neuroscience 2019;22:1371–1378 doi: 10.1038/s41593-019-0436-x.

6. Bürk K. Ataxia Scales for the Clinical Evaluation. In: Gruol DL, Koibuchi N, Manto M, Molinari M, Schmahmann JD, Shen Y, editors. Essentials of Cerebellum and Cerebellar Disorders: A Primer For Graduate Students. Cham: Springer International Publishing; 2016. pp. 513–520. doi: 10.1007/978-3-319-24551-5_69.

7. Fatemi SH. Cerebellar Pathology in Autism. In: Gruol DL, Koibuchi N, Manto M, Molinari M, Schmahmann JD, Shen Y, editors. Essentials of Cerebellum and Cerebellar Disorders: A Primer For Graduate Students. Cham: Springer International Publishing; 2016. pp. 539–543. doi: 10.1007/978-3-319-24551-5_72.

8. Kheradmand A, Kim JS, Zee D. Cerebellum and Oculomotor Deficits. In: Gruol DL, Koibuchi N, Manto M, Molinari M, Schmahmann JD, Shen Y, editors. Essentials of Cerebellum and Cerebellar Disorders: A Primer For Graduate Students. Cham: Springer International Publishing; 2016. pp. 471–475. doi: 10.1007/978-3-319-24551-5_64.

9. Schmahmann JD. The Cerebellar Cognitive Affective Syndrome and the Neuropsychiatry of the Cerebellum. In: Gruol DL, Koibuchi N, Manto M, Molinari M, Schmahmann JD, Shen Y, editors. Essentials of Cerebellum and Cerebellar Disorders: A Primer For Graduate Students. Cham: Springer International Publishing; 2016. pp. 499–511. doi: 10.1007/978-3-319-24551-5_68.

10. Oz G, Alger JR, Barker PB, et al. Clinical proton MR spectroscopy in central nervous system disorders. Radiology 2014;270:658–679 doi: 10.1148/radiol.13130531.

11. Deelchand DK, Adanyeguh IM, Emir UE, et al. Two-site reproducibility of cerebellar and brainstem neurochemical profiles with short-echo, single-voxel MRS at 3T. Magnetic resonance in medicine 2015;73:1718–1725.

12. Long Z, Dyke JP, Ma R, Huang CC, Louis ED, Dydak U. Reproducibility and effect of tissue composition on cerebellar GABA MRS in an elderly population. NMR Biomed 2015;28:1315–1323 doi: 10.1002/nbm.3381.

13. Maudsley AA, Domenig C, Govind V, et al. Mapping of Brain Metabolite Distributions by Volumetric Proton MR Spectroscopic Imaging (MRSI). Magn Reson Med 2009;61:548–559 doi: 10.1002/mrm.21875.

14. Jacobs MA, Horská A, van Zijl PC, Barker PB. Quantitative proton MR spectroscopic imaging of normal human cerebellum and brain stem. Magn Reson Med 2001;46:699–705.

15. Pfeuffer J, van de Moortele P-F, Yacoub E, et al. Zoomed Functional Imaging in the Human Brain at 7 Tesla with Simultaneous High Spatial and High Temporal Resolution. NeuroImage 2002;17:272–286 doi: 10.1006/nimg.2002.1103.

16. Sun H, Fessler JA, Noll DC, Nielsen J-F. Rapid inner-volume imaging in the steady-state with 3D selective excitation and small-tip fast recovery (STFR) imaging. Magn Reson Med 2016;76:1217–1223 doi: 10.1002/mrm.26026.

17. Golay X, Gillen J, van Zijl PCM, Barker PB. Scan time reduction in proton magnetic resonance spectroscopic imaging of the human brain. Magn Reson Med 2002;47:384–387 doi: 10.1002/mrm.10038.

18. Maudsley AA, Matson GB, Hugg JW, Weiner MW. Reduced phase encoding in spectroscopic imaging. Magnetic Resonance in Medicine 1994;31:645–651 doi: 10.1002/mrm.1910310610.

19. Emir UE, Burns B, Chiew M, Jezzard P, Thomas MA. Non-water-suppressed short-echo-time magnetic resonance spectroscopic imaging using a concentric ring k-space trajectory. NMR Biomed 2017;30 doi: 10.1002/nbm.3714.

20. Chiew M, Jiang W, Burns B, et al. Density-weighted concentric rings k-space trajectory for 1H magnetic resonance spectroscopic imaging at 7 T. NMR Biomed 2018;31 doi: 10.1002/nbm.3838.

21. Steel A, Chiew M, Jezzard P, et al. Metabolite-cycled density-weighted concentric rings k-space trajectory (DW-CRT) enables high-resolution 1LH magnetic resonance spectroscopic imaging at 3-Tesla. Scientific Reports 2018;8:7792 doi: 10.1038/s41598-018-26096-y.

22. Fessler JA, Sutton BP. Nonuniform fast Fourier transforms using min-max interpolation. IEEE Transactions on Signal Processing 2003;51:560–574 doi: 10.1109/TSP.2002.807005.

23. Cabanes E, Confort-Gouny S, Le Fur Y, Simond G, Cozzone PJ. Optimization of residual water signal removal by HLSVD on simulated short echo time proton MR spectra of the human brain. J. Magn. Reson. 2001;150:116–125 doi: 10.1006/jmre.2001.2318.

24. Bilgic B, Chatnuntawech I, Fan AP, et al. Fast image reconstruction with L2-regularization. J Magn Reson Imaging 2014;40:181–191 doi: 10.1002/jmri.24365.

25. Provencher SW. Automatic quantitation of localized in vivo 1H spectra with LCModel. NMR Biomed 2001;14:260–264.

26. Quadrelli S, Mountford C, Ramadan S. Hitchhiker’s Guide to Voxel Segmentation for Partial Volume Correction of In Vivo Magnetic Resonance Spectroscopy. Magn Reson Insights 2016;9:1–8 doi: 10.4137/MRI.S32903.

27. Jenkinson M, Beckmann CF, Behrens TEJ, Woolrich MW, Smith SM. FSL. NeuroImage 2012;62:782–790 doi: 10.1016/j.neuroimage.2011.09.015.

28. Benjamini Y, Hochberg Y. Controlling the False Discovery Rate: A Practical and Powerful Approach to Multiple Testing. Journal of the Royal Statistical Society. Series B (Methodological) 1995;57:289–300.

29. Alhulail AA, Patterson DA, Xia P, et al. Fat–water separation by fast metabolite cycling magnetic resonance spectroscopic imaging at 3 T: A method to generate separate quantitative distribution maps of musculoskeletal lipid components. Magnetic Resonance in Medicine n/a doi: 10.1002/mrm.28228.

30. Keil B, Blau JN, Biber S, et al. A 64-channel 3T array coil for accelerated brain MRI. Magn Reson Med 2013;70:248–258 doi: 10.1002/mrm.24427.

31. Chang P, Nassirpour S, Avdievitch N, Henning A. Non-water-suppressed 1 H FID-MRSI at 3T and 9.4T. Magn Reson Med 2018;80:442–451 doi: 10.1002/mrm.27049.

32. Zhang SH, Maier SE, Panych LP. Improved spatial localization in magnetic resonance spectroscopic imaging with two-dimensional PSF-Choice encoding. J. Magn. Reson. 2018;290:18–28 doi: 10.1016/j.jmr.2018.03.002.

33. Klauser A, Courvoisier S, Kasten J, et al. Fast high-resolution brain metabolite mapping on a clinical 3T MRI by accelerate H-FID-MRSI and low-rank constrained reconstruction. Magn Reson Med 2019;81:2841–2857 doi: 10.1002/mrm.27623.

34. Ma C, Lam F, Ning Q, Johnson CL, Liang Z-P. High-resolution 1 H-MRSI of the brain using short-TE SPICE. Magn Reson Med 2017;77:467–479 doi: 10.1002/mrm.26130.

35. Hangel G, Strasser B, Považan M, et al. Ultra-high resolution brain metabolite mapping at 7 T by short-TR Hadamard-encoded FID-MRSI. Neuroimage 2018;168:199–210 doi: 10.1016/j.neuroimage.2016.10.043.

36. Emir UE, Auerbach EJ, Moortele P-FVD, et al. Regional neurochemical profiles in the human brain measured by 1H MRS at 7 T using local B1 shimming. NMR in Biomedicine 2012;25:152–160.

37. Jaeger D, Lu H. Cerebellar Nuclei. In: Gruol DL, Koibuchi N, Manto M, Molinari M, Schmahmann JD, Shen Y, editors. Essentials of Cerebellum and Cerebellar Disorders: A Primer For Graduate Students. Cham: Springer International Publishing; 2016. pp. 311–315. doi: 10.1007/978-3-319-24551-5_42.

38. Moffett JR, Namboodiri AMA. Expression of N-Acetylaspartate and N-Acetylaspartylglutamate in the Nervous System. In: Moffett JR, Tieman SB, Weinberger DR, Coyle JT, Namboodiri AMA, editors. N-Acetylaspartate. Advances in Experimental Medicine and Biology. Boston, MA: Springer US; 2006. pp. 7–26. doi: 10.1007/0-387-30172-0_2.

39. Fa A, Lr C, Lt G, et al. Equal Numbers of Neuronal and Nonneuronal Cells Make the Human Brain an Isometrically Scaled-Up Primate Brain. The Journal of comparative neurology. https://pubmed.ncbi.nlm.nih.gov/19226510/?dopt=Abstract. Published April 10, 2009. Accessed May 19, 2020 doi: 10.1002/cne.21974.

40. Palomero-Gallagher N, Zilles K. Chapter 24 - Cyto- and receptor architectonic mapping of the human brain. In: Huitinga I, Webster MJ, editors. Handbook of Clinical Neurology. Vol. 150. Brain Banking. Elsevier; 2018. pp. 355–387. doi: 10.1016/B978-0-444-63639-3.00024-4.

41. Lecocq A, Le Fur Y, Maudsley AA, et al. Whole-brain quantitative mapping of metabolites using short echo 3D-proton-MRSI. J Magn Reson Imaging 2015;42:280–289 doi: 10.1002/jmri.24809.

42. Tannús A, Garwood M. Adiabatic pulses. NMR Biomed 1997;10:423–434.

